# Use of Induced Pluripotent Stem Cells to Build Isogenic Systems and Investigate Type 1 Diabetes

**DOI:** 10.1101/2021.05.03.441837

**Authors:** Lucas H. Armitage, Scott E. Stimpson, Katherine E. Santostefano, Lina Sui, Similoluwa O. Ogundare, Brittney N. Newby, Roberto Castro-Gutierrez, Mollie K. Huber, Jared P. Taylor, Prerana Sharma, Ilian Radichev, Daniel J. Perry, Natalie C. Fredette, Alexei Savinov, Mark A. Wallet, Naohiro Terada, Todd M. Brusko, Holger A. Russ, Jing Chen, Dieter Egli, Clayton. E Mathews

## Abstract

Type 1 diabetes is a disease that arises due to complex immunogenetic mechanisms. Key cell-cell interactions involved in the pathogenesis of T1D are activation of autoreactive T cells by dendritic cells (DC), migration of T cells across endothelial cells (EC) lining capillary walls into the islets of Langerhans, interaction of T cells with macrophages in the islets, and killing of β-cells by autoreactive CD8^+^ T cells. Overall, pathogenic cell-cell interactions are likely regulated by the individual’s collection of genetic T1D-risk variants. To accurately model the role of genetics, it is essential to build systems to interrogate single candidate genes in isolation during the interactions of cells that are essential for disease development. However, obtaining single-donor matched cells relevant to T1D is a challenge. Sourcing these genetic variants from human induced pluripotent stem cells (iPSC) avoids this limitation. Herein, we have differentiated iPSC from one donor into DC, macrophages, EC, and β-cells. Additionally, we also engineered T cell avatars from the same donor to provide an *in vitro* platform to study genetic influences on these critical cellular interactions. This proof of concept demonstrates the ability to derive an isogenic system from a single donor to study these relevant cell-cell interactions. Our system constitutes an interdisciplinary approach with a controlled environment that provides a proof-of-concept for future studies to determine the role of disease alleles (e.g. *IFIH1, PTPN22, SH2B3, TYK2)* in regulating cell-cell interactions and cell-specific contributions to the pathogenesis of T1D.

## 1 Introduction

Type 1 diabetes (T1D) is a debilitating autoimmune disease that is caused by T cell-mediated destruction of β-cells in the islets of Langerhans in the pancreas. This results in lifelong dependence on exogenous insulin and can lead to many complications that degrade quality of life for patients living with T1D. With the advent of genome-wide association studies (GWAS) a big picture view of the genetic contributions of T1D has come into focus. While human leukocyte antigen (HLA) loci, encoding the major histocompatibility (MHC) proteins, make up the bulk of genetic risk for T1D, over 100 non-HLA loci have been identified with polymorphisms that increase or decrease risk for T1D (1,2,11,12,3–10). However, studying the phenotypic outcome of these polymorphisms has proven difficult due to the heterogeneous nature of the human population and the inaccessibility of pancreatic tissue from individuals with signs of islet autoimmunity (i.e. islet autoantibodies) and individuals with clinical T1D.

Inbred rodent models, such as the non-obese diabetic (NOD) mouse and the Biobreeding rat, have been paramount in the parsing out of key cellular interactions that lead to the initiation of immune dysfunction and eventually autoimmune diabetes. Indeed, a number of the genes or regions mapped that contribute to the development of diabetes in the NOD mouse are orthologous, syntenic, or impact the same pathways as genes/loci contributing to human T1D risk (13). However, due to species dissimilarity, many genes that increase or decrease risk for T1D in humans are not representative of T1D-risk in NOD mice. For example, the single nucleotide polymorphism (SNP) *rs2476601* in *PTPN22* has no analogous polymorphism in NOD mice. Both the major allele of *PTPN22* and the mouse orthologue, *Ptpn22*, are negative regulators of T cell receptor (TCR) and B cell receptor (BCR) signaling in humans and mice. However, the arginine 620 to tryptophan conversion (R620W) encoded by the minor allele at *rs2476601* has disparate effects when introduced to the mouse genome. In humans, the minor allele of *PTPN22* blunts while in mice this amino acid substitution potentiates BCR signaling (14–17). With the advent of efficient gene editing technologies and methods to differentiate induced pluripotent stem cell (iPSC) clones to defined cell types, our goal is to develop an isogenic system, derived from a renewable cell source, that allows for interrogation of the pathogenic responsibility of single T1D-linked alleles as well as the cells where they are expressed to explore human cell function as well as cell-cell interactions that influence T1D.

Over the past half-century, human and murine systems have been implemented and both immune and non-immune cells have been implicated in T1D (18). Key interactions involved in the pathogenesis of T1D in humans (Figure 1) include; 1) the initial interactions between dendritic cells (DC) and β-cells where DC uptake β-cell antigens from dead, dying, or stressed β-cells, 2) the activation of autoreactive CD4^+^ and CD8^+^ T cells by the β-cell antigen-loaded DC, 3) the interaction between autoreactive CD8^+^ T cells and the endothelial cells (EC) lining the capillaries that supply the islets of Langerhans as autoreactive CD8^+^ T cells undergo transendothelial migration to access the islets, 4.1) the targeting of β-cells for cell-mediated lysis by autoreactive CD8^+^ T cells, and 4.2) the *in situ* amplification of T cell responses by DC and macrophages in the islets. To develop an isogenic model there is a need to isolate cell types of interest from a single donor, differentiate cell types from donor stem cells, or use a combination of these two approaches. The utility of such single-donor iPSCs was recently highlighted in a study that exemplifies the relationship between rare and common gene variants and gene expression in iPSCs (19).

**Figure 1.**
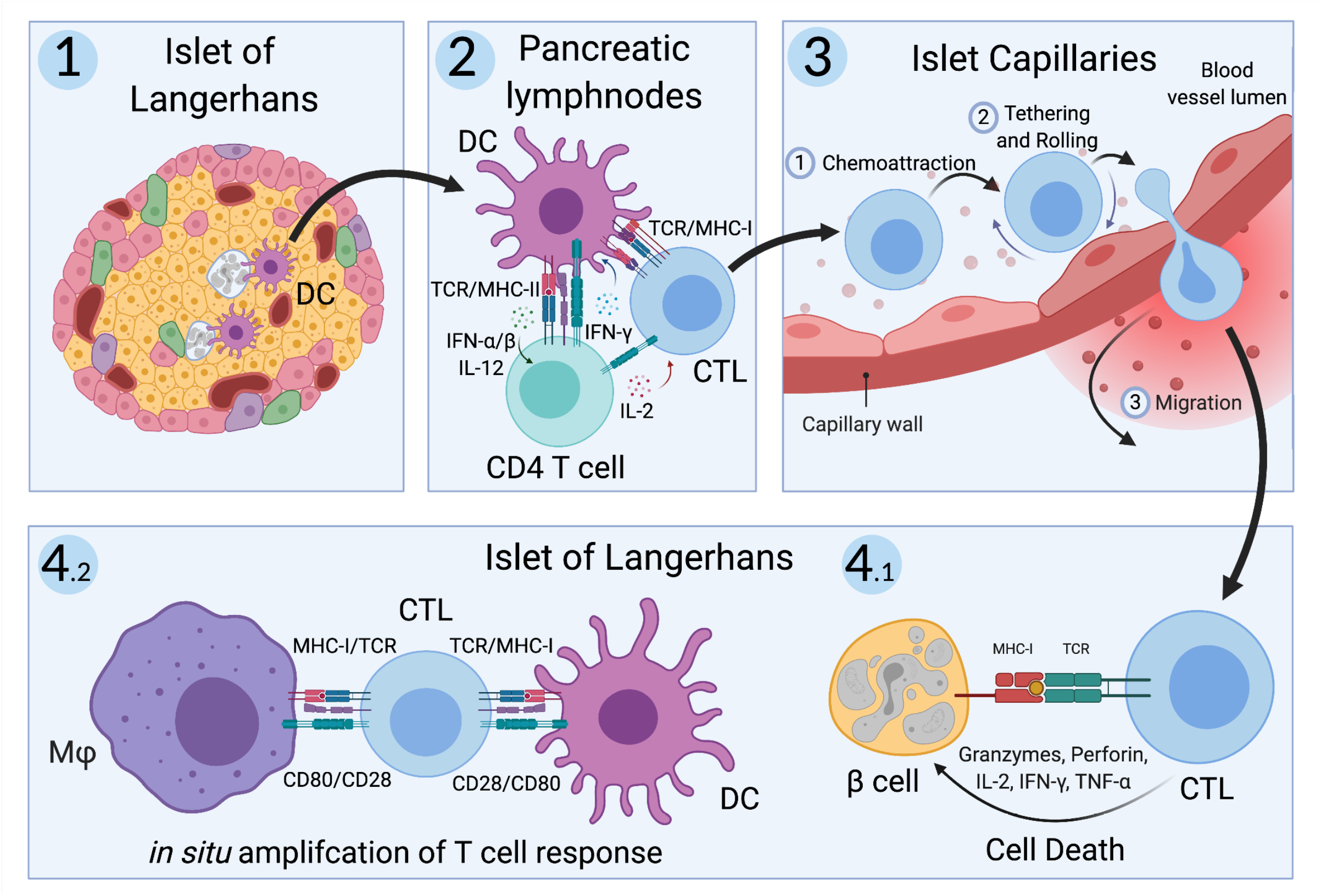
Key cellular interactions involved in pathogenesis of T1D in humans. First, dendritic cells (DC) uptake antigen from dead, dying, and stressed β-cells in the islets of Langerhans in individuals at risk for type 1 diabetes (T1D). Second, these β-cell-antigen-loaded DC migrate to the pancreatic lymph nodes where they present β-cell antigen to autoreactive CD4^+^ and CD8^+^ T cells, initiating an adaptive immune response. Third, these autoreactive CD8^+^ T cells, also called cytotoxic T lymphocytes (CTLs), migrate back to the pancreas and undergo transendothelial migration across the endothelium lining capillaries that supply the islets of Langerhans. Fourth, these CTLs directly target and lyse β-cells while resident and infiltrating macrophages and DC amplify the CTL response *in situ*. Created with BioRender.com.

Human iPSC share similar gene expression patterns to human embryonic stem cells (ESC) and have the capacity to differentiate to all cell types of the human body. Generating iPSC and differentiating them to the cell types of interest from a single donor preserves the genetic background and integrity of a particular disease allele potentially making for an excellent isogenic model system to study T1D pathogenesis. In support of these isogenic systems, advances in stem cell technologies have resulted in protocols allowing for differentiation of iPSC to functional replicas of cells essential for T1D, including hard to access cells, i.e. β-cells. Additionally, methods for engineering primary human T cells have advanced our understanding of immunity and autoimmunity (20–22). Here we selected 3 candidate iPSC lines to build the isogenic system, 1-018, 1-023, and 2395, that had previously been differentiated to stem cell derived beta-like cells (sBC), as this is the most time intensive and thus the rate limiting step for construction of these isogenic models (23–25). Based on HLA, we narrowed down our selection by eliminating 1-018 as our CD8^+^ T cell receptors (TCRs) are all HLA-A*0201 restricted and 1-018 is HLA-A*0101/0301. As we worked to differentiate the iPSC lines to the other cells of interest, we were unable to induce monocyte production from line 1-023 and focused solely on 2395. We have successfully differentiated 2395 iPSC into pancreatic β-cells, endothelial cells, macrophages, and dendritic cells. In addition, T cells were purified from this same donor. Employing the most recent and advanced methods for culturing and differentiating iPSC that have similar functionality to cells *in vivo*, we show that these iPSC-derived cells can be used for modeling and interrogating cell-cell interactions that are relevant to the pathogenesis of T1D (Figure 1). Utilizing these cellular approaches will allow investigators to interrogate relationships of genotype with cell function and cell-cell interactions.

## 2 Materials and Methods

### 2.1 Reprogramming somatic cells to iPSC

The generation and characterization of the iPSC lines 1-018 and 1-023 was previously described (23–25). Derivation of iPSC from donor 2395 was as previously described (26). Briefly, peripheral blood was collected from donor 2395 under informed consent by the University of Florida Diabetes Institute Study Bank with Institutional Review Board approval (IRB201400703). CD34^+^ peripheral stem cells were isolated from peripheral blood and expanded using the Complete Kit for Human Whole Blood CD34^+^ Cells (Stem Cell Technologies, Vancouver, Canada) following the manufacturer’s protocol. Expanded CD34^+^ peripheral stem cells were transduced with the Sendai viral vector, SeVdp(KOSM)302L, which encodes the 4 reprogramming factors, OCT4, SOX2, KLF4, and c-MYC (27, 28). Following transduction, cells were cultured on hESC qualified Matrigel- (Corning, Corning, NY) coated dishes in ReproTeSR medium (Stem Cell Technologies) and medium was changed daily until iPSC colonies were observed. iPSC colonies were manually isolated then expanded on Matrigel-coated dishes in mTeSR1 medium (StemCell Technologies, Vancouver, Canada).

Following reprogramming, donor 2395 iPSC were assessed for pluripotency using the StemDiff Trilineage Differentiation Kit (StemCell Technologies) following the manufacturer’s protocol. iPSC were subjected to karyotyping to verify a normal karyotype by Cell Line Genetics (Madison, WI, USA). Undifferentiated donor 2395 iPSC were stained with anti-SSEA-4-APC and anti-Oct-4-PE (BioLegend, San Diego, CA) using the Fixation/Permeabilization Solution kit from BD Biosciences (San Jose, CA) and analyzed on an Accuri C6 flow cytometer (BD Biosciences). Information for all antibodies used during the included studies are contained in the Supplemental Table.

### 2.2 Differentiation of iPSC to Stem Cell-Derived beta-like Cells

The iPSC lines used in this study, 1-018, 1-023, and 2395, were initially selected because they had previously been differentiated to sBC. The iPSC lines, 1-018 and 1-023, were differentiated to sBC as previously described (25). Donor 2395 iPSC were maintained on Matrigel-coated 6 well plates in mTeSR+ medium (StemCell Technologies) and differentiated to sBC as previously described with some modifications (29). iPSC were maintained on Matrigel-coated 6 well plates in mTeSR+ medium (StemCell Technologies). Differentiation to sBC was carried out in suspension□based, low attachment suspension culture plates as described (30) or in an ABLE bioreactor magnetic stirring system (Reprocell, Beltsville, MD, USA) as follows. For differentiations, 70 - 80% confluent cultures were washed with PBS and incubated in TrypLE Express Enzyme (Gibco, Watham, MD, USA) for 8 min at 37°C followed by quenching with mTeSR+. Bioreactors were seeded at 0.5 × 10^6^ cells/mL in mTeSR+ medium supplemented with 10 mM ROCK inhibitor. Bioreactors were placed on a magnetic stirring system set at 60 RPM in a cell culture incubator at 5% CO_2_ to induce sphere formation for 48-72 hours. To induce definitive endoderm differentiation, spheres were collected in a 50 mL Falcon tube, allowed to settle by gravity, washed once with RPMI (Gibco) + 0.2 % FBS, and re□suspended in day 0 medium (RPMI containing 0.2 % FBS, 1:5,000 ITS (Gibco), 100 ng/mL Activin-A (R&D Systems), and 3 mM CHIR (StemCell Technologies)). The differentiation medium was changed daily by letting spheres settle by gravity for 3-10 min. Approximately 80% of spent supernatant was removed by aspiration, fresh medium was added, and bioreactors were placed back on stirrer system. The sBC differentiation was based on a previously published protocol (29) with modifications as outlined below. Differentiation media are as follows: day 1 and 2, RPMI containing 0.2 % FBS, 1:2,000 ITS, and 100 ng/mL Activin A; day 3 and 4, RPMI containing 2% FBS, 1:1,000 ITS, and 50 ng/LmL KGF (Peprotech); day 5, DMEM with 4.5 g/L D-glucose (Gibco) containing 1:100 SM1 (StemCell Technologies), 1:100 NEAA (Gibco), 1 mM Sodium Pyruvate (Gibco), 1:100 GlutaMAX (Gibco), 3 nM TTNPB, (R&D Systems), 250 nM Sant-1 (R&D Systems), 250 nM LDN (StemCell Technologies), 30 nM PMA (Sigma Aldrich), 50 mg/mL 2-phospho-L-ascorbic acid trisodium salt (VitC; Sigma); day 6, DMEM with 4.5 g/L D-glucose containing 1:100 SM1, 1:100 NEAA, 1 mM Sodium Pyruvate, 1:100 GlutaMAX, 3 nM TTNPB and 50 mg/mL VitC; day 7, DMEM containing 1:100 SM1, 1:100 NEAA, 1 mM Sodium Pyruvate, 1:100 GlutaMAX, 3 nM TTNPB and 50 mg/mL VitC; day 8 and 9, DMEM containing 1:100 SM1, 1:100 NEAA, 1 mM Sodium Pyruvate, 1:100 GlutaMAX, 200 ng/ml EGF and 50 ng/mL KGF; days 10-16, DMEM containing 2 % fraction V BSA, 1:100 NEAA, 1 mM Sodium Pyruvate, 1:100 GlutaMAX, 1:100 ITS, 10 mg/ml Heparin (Sigma), 2 mM N-Acetyl-L-cysteine (Cysteine; Sigma), 10 mM Zinc sulfate heptahydrate (Zinc; Sigma), 1x BME, 10 mM Alk5i II RepSox (R&D Systems), 1 mM 3,3’,5-Triiodo-L-thyronine sodium salt (T3) (Sigma), 0.5 mM LDN, 1 mM Gamma Secretase Inhibitor XX (XXi) (AsisChem, Waltham, MA, USA) and 1:250 1 M NaOH to adjust pH to ∼7.4; day 17 and up, CMRL (Gibco) containing 1% BSA, 1:100 NEAA, 1 mM Sodium Pyruvate, 1:100 GlutaMAX, 10 mg/mL Heparin, 2 mM Cysteine, 10 mM Zinc, 1x BME, 10 mM Alk5i II RepSox, 1 mM T3, 50 mg/mL VitC, and 1:250 NaOH to adjust pH to ∼7.4 (also referred to as maturation medium). All media contained 1x PenStrep (Gibco). Medium was changed every other day starting day 11. Validation and function of sBC, including staining for glucagon and insulin as well as glucose-stimulated insulin secretion were performed as previously described (25,29,30).

### 2.3 HLA Typing of iPSC

DNA was isolated from iPSC lines, 1-018, 1-023, and 2395, and subjected to genotyping at 974,650 unique loci using a custom SNP array. The Axiom Precision Medicine Research Array (ThermoFisher Scientific) was modified to include content from the ImmunoChip v2.0 (31) as well as previously reported credible T1D risk variants (2), and additional content to create the UFDIchip. UFDIchips were processed on an Affymetrix Gene Titan instrument with external sample handling on a BioMek FX dual arm robotic workstation. Data processing and quality control pipelines included standard quality control procedures at the SNP, sample, and plate levels using Axiom™ Analysis Suite 3.0 (ThermoFisher Scientific) set to the default stringency thresholds as recommended, as well as for genetic versus reported sex. The analysis pipeline also includes race imputation using EthSeq (32), relatedness using KING (33), and imputation to 40M SNPs using the Human Reference Consortium (version 1.1) with the Michigan Imputation Server (34). The array also includes dense coverage of the highly polymorphic HLA region allowing accurate imputation of HLA haplotypes to 4-digit resolution (35).

### 2.4 Differentiation of iPSC to Monocytes, Monocyte-derived Macrophages, and Monocyte-derived Dendritic Cells

Donor 2395 iPSC were differentiated to monocytes, monocyte-derived dendritic cells (moDC, and monocyte-derived macrophages (MDM), as described (36). Briefly, iPSC were differentiated to hematopoietic progenitors using the STEMdiff Hematopoietic Kit (StemCell Technologies) over 12 days. On day 12, hematopoietic progenitors were harvested and the hemogenic endothelium left behind in the 12-well plate was cultured in X-VIVO15 (Lonza, Basel, Switzerland) supplemented with β-mercaptoethanol (Gibco), Glutamax (Gibco), 25 ng/mL recombinant human IL-3 (rhIL-3, PeproTech, Rocky Hill, NJ), and 100 ng/mL recombinant human M-CSF (rhM-CSF, PeproTech). The hemogenic endothelium transitioned into monocyte factories and produced non-adherent CD14^+^ monocytes that can be harvested repeatedly starting around day 21. Thereafter, CD14^+^ cells are harvested every 2-7 days. Monocyte factories were replaced on Day 94. iPSC-monocytes were differentiated to iPSC-MDM as previously described (26). Briefly, iPSC-monocytes were plated in high-glucose DMEM supplemented with 10% FBS (Genesee Scientific, San Diego, CA) or 10% human serum (Atlanta Biologicals, Flowery Branch, GA), 1000 U/mL Penicillin/streptomycin (Genessee Scientific), Glutamax, and 10 ng/ml of rhM-CSF for 7 days before use in experiments. iPSC-monocytes from Donor 2395 were differentiated to iPSC-moDC as previously described (37). Briefly, the CD14^+^ iPSC-monocytes were plated in GMP DC medium (CellGenix, Freiburg im Breisgau, Germany) supplemented with Glutamax, 50 ng/mL recombinant human IL-4 (PeproTech), and 50 ng/mL recombinant human GM-CSF (PeproTech) and cultured for 6 days. On day 7, iPSC-moDC were plated for experiments.

### 2.5 Differentiation of iPSC to Endothelial Cells

Donor 2395 iPSC were differentiated to endothelial cells (iEC) as previously described (38). This newly published protocol was adapted from a previously published protocol (39). Briefly, donor 2395 iPSC colonies were dissociated to single cells with Accutase (StemCell Technologies) and plated on Matrigel-coated plates at a density of 5 x 10^5^ cells per cm^2^ in mTESR1 medium (StemCell Technologies) supplemented with 10 ng/mL of the RHO/ROCK pathway inhibitor, Y-27632 (StemCell Technologies) on day 0. On day 1, the medium was changed to STEMdiff Mesoderm Induction medium (StemCell Technologies) and the medium was changed daily. On day 3, medium was replaced with STEMdiff APEL medium (StemCell Technologies) supplemented with 25 ng/mL BMP-4 (PeproTech) and 50 ng/mL VEGF (Peprotech). On day 5, the medium was replaced with fresh STEMdiff APEL medium supplemented with 25 ng/mL BMP-4 and 50 ng/mL VEGF. On day 7, CD34^+^ cells were enriched with the EasySep Human Cord Blood CD34 Positive Selection Kit II (StemCell Technologies) without using the RosetteSep™ Human Cord Blood CD34 Pre-Enrichment Cocktail. The CD34^+^ cells were plated at a density of 3.5 x 10^5^ cells per cm^2^ onto fibronectin (Millipore) coated dishes (5 ng/cm^2^) and cultured in Endothelial Growth Medium 2 (EGM2, Lonza) supplemented with 50 ng/mL VEGF with medium changes every other day until they reached confluency. After the 1st passage, cells were cultured in regular EGM2 without extra VEGF and split at a 1:3 ratio onto fibronectin-coated plates (5 ng/cm^2^) as they approached confluency. iEC were cryopreserved in 0.5mL of CryoStor CS10 Freeze Media (BioLife Solutions, Bothell, WA) at 1 x 10^6^ cells/vial according to the manufacturer’s protocol.

### 2.6 Generation of CD8^+^ T Cell Avatars from Donor 2395

Antigen-specific CD8^+^ T Cell avatars were produced and validated as previously published (20). Briefly, CD8^+^ T cells were isolated from peripheral blood of donor 2395 via negative selection using the RosetteSep Human CD8^+^ T Cell Enrichment Cocktail (StemCell Technologies) according to the manufacturer’s protocol. Naïve CD8^+^ T cells (CD45RA^+^CD45RO^-^) were isolated by FACS with a FACSAria III Cell Sorter (BD Biosciences). Naïve CD8^+^ T cells were activated for two days with Human T-Activator CD3/CD28 Dynabeads (ThermoFisher Scientific, Waltham, MA). After two days, cells were transduced with lentiviral vectors, either pCCL.IGRPopt.eGFP, encoding a TCR recognizing an HLA-A*02-01-restricted epitope derived from the T1D-relevant autoantigen, glucose-6-phosphatase 2 (G6PC2) commonly referred to as islet-specific G6CP-related protein (IGRP) (40, 41) or LV.Mart1.TCR.RK, encoding a TCR recognizing an HLA-A*02-01-restricted epitope derived from the non-T1D-relevant auto-antigen, MART-1 (21, 22). Following transduction, the IGRP and the MART-1 CD8+ T cell avatars (IGRP avatars and MART-1 avatars) were expanded for 7 more days then cryopreserved.

### 2.7 Co-culturing Donor 2395 iPSC-moDC and iPSC-MDM with Donor 2395 CD8^+^ T Cells

To verify the ability of iPSC-moDC and iPSC-MDM to induce antigen-specific T cell expansion, iPSC-moDC and MDM were treated with 10 ng/mL of LPS (Sigma-Aldrich, St. Louis, MO) and 10 ng/mL of IFNγ (Peprotech) for 48 hours. After iPSC-moDC and MDM were activated for 48 hours, they were loaded for 2 hours with both CEFX Ultra SuperStim Pool MHC-I and MHC-II Subsets (JPT innovative peptide solutions, Berlin, Germany), for a total of 80 MHC Class I restricted epitopes and 68 MHC Class II restricted epitopes derived from a broad range of common human viruses, as well as *Clostridium tetani* and *Toxoplasma gondii*. After 2 hours, iPSC-moDC and MDM were washed with T cell expansion medium (RPMI supplemented with 10% HyClone FBS (GE Healthcare, Chicago, IL, USA), 1000 U/mL Penicillin/streptomycin, 1mM Sodium Pyruvate (Corning), 10mM HEPES (Corning), nonessential amino acids (Corning), Glutamax, and β-mercaptoethanol) and the medium was replaced with T cell expansion medium. Freshly isolated total T cells from donor 2395 were stained with CellTrace CFSE (ThermoFisher), added to the CEFX peptide-loaded iPSC-moDC or MDM and co-cultured for 5 days. After 5 days, expansion of total T cells was analyzed via flow cytometry on an Accuri C6 Flow Cytometer.

### 2.8 *In Vitro* Hydrodynamic Flow Chamber Adhesion Assay

The *in vitro* hydrodynamic flow chamber adhesion assay was performed as previously described (42, 43). Briefly, iEC were grown in monolayers to 95-100% confluence on a 0.2 mm Luer µ-slide (channel slide) pre-coated with fibronectin. The iEC monolayers were then loaded with either vehicle solution, IGRP_265-273_ (the antigenic peptide recognized by the IGRP avatars) or Melan-A_26-35_ (the antigenic peptide recognized by the MART-1 avatars) at a final concentration of 10 µg/ml and incubated overnight. The next day, the iEC-containing channel slide(s) were loaded on the stage of a confocal microscope heated to 37°C. Ten thousand MART-1 or IGRP avatars labeled with 200 nM CellTracker Green (Molecular Probes, Eugene, OR) were added to the channel slide and allowed to settle for 30 minutes. HBSS buffer with MgCl_2_ and CaCl_2_ was then flowed across the channel slide by a syringe-driven pump at a series of increasing flow rates ranging from 1 to 80 dyne/cm^2^. After passing the buffer through the channel slide for 1 minute for each shear flow rate, bright field and fluorescent images of the slide were acquired by confocal microscopy. T cells that remained attached after each flow rate were then counted using ImageJ software (44).

### 2.9 Chromium Release Assay

The chromium release assay to assess antigen-specific targeting of sBC or the immortalized human β-cell line, BetaLox5 cells (βL5), by donor 2395 avatars was performed as previously described (20, 45). Briefly, sBC clusters from donor 2395, 1-018, or 1-023 were dispersed in enzyme-free Cell Dissociation Buffer (Gibco) for 10 minutes at 37°C with gentle pipetting. Dispersed sBC were plated at 40,000 cells/well or βL5 cells were plated at 10,000 cells/well in 96-well flat-bottom plates. Target cells were loaded with Melan-A_27-35_. Target cells were then labeled with ^51^Cr at 1 µCi/well for 3 hours and then washed 3 times with culture medium. Donor 2395 MART-1 avatars were added at effector to target (E:T) ratios of 0:1, 5:1, and 10:1 and co-cultured for 16 hours. Afterwards, the supernatants were collected, cells were lysed in 2% SDS, and cell lysate was collected. ^51^Cr release was measured with a gamma counter and specific lysis was calculated as follows: 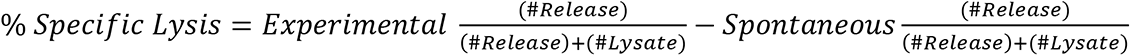 where spontaneous release is the 0:1 E:T ratio and experimental release is measured in the 5:1 and 10:1 E:T ratios.

### 2.10 Statistical Analysis

All statistical analyses were performed in GraphPad Prism version 8.4.3 (San Diego, CA) using unpaired Student’s t test, one-way ANOVA, or two-way ANOVA, as indicated in the figure legends.

## 3 Results

### 3.1 Reprogramming Donor 2395 CD34^+^ Peripheral Stem Cells to iPSC

Our proposed isogenic system for T1D (Figure 1) requires constant supplies of β-cells, dendritic cells, T cells, endothelial cells, and macrophages from the same donor. The major goal of this effort was to determine if iPSC lines had potential to differentiate into all of these cell types allowing construction of a fully isogenic system. For the work described here all of these cell types, except for T cells, were to be differentiated from donor iPSC lines. Before working with these iPSC lines, they were all verified to express pluripotency markers and have a normal karyotype. iPSC from lines 1-023 and 1-018 were validated and published previously (23–25). iPSC generated from donor 2395 showed similar morphology to normal hESCs (Figure 2A) and expressed the pluripotency markers, Oct4 and SSEA-4 (Figure 2B). 2395 iPSC were pluripotent, they differentiated into Nestin^+^ ectoderm, FoxA2^+^Sox17^+^ endoderm, and NCAM^+^Brachyury^+^ mesoderm (Figure 2C). Karyotyping analysis of 2395 iPSC showed a normal chromosome count of 46,XY (Figure 2D). Furthermore, 2395 iPSC have been passaged more than 35 times without any reduction in cell proliferation.

**Figure 2.**
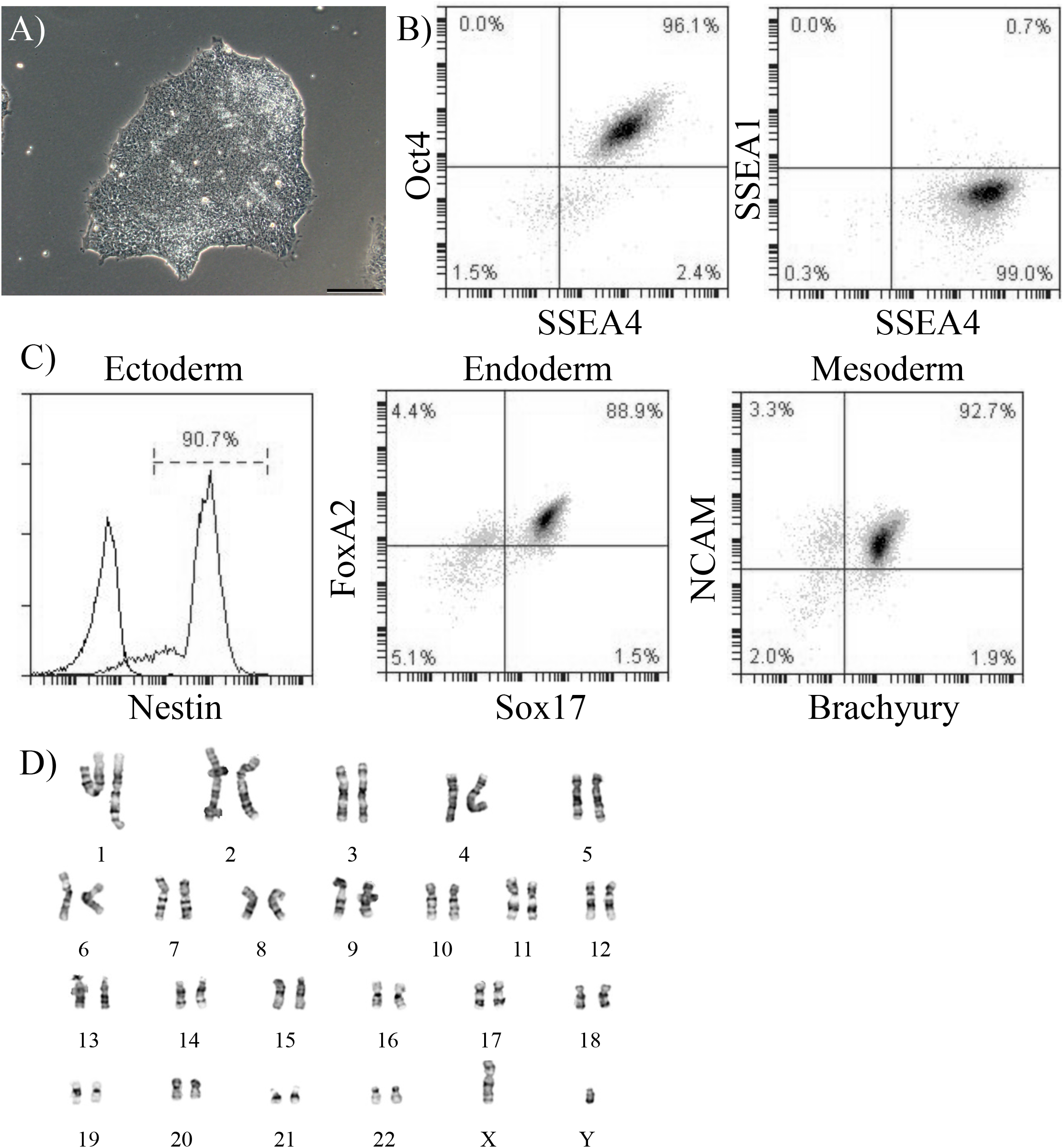
Verification of pluripotency of donor 2395 iPSC. **(A)** A phase contrast image of an iPSC colony from donor 2395 showing densely packed cells with a defined border characteristic of iPSC. **(B)** Donor 2395 iPSC are positive for Oct4 and SSEA4 while negative for SSEA1 expression. **(C)** Donor 2395 iPSC express Nestin when differentiated to ectoderm, express FoxA2 and Sox17 when differentiated to endoderm, and express NCAM and Brachyury when differentiated to mesoderm. **(D)** Karyotypic analysis of donor 2395 iPSC reveals a normal karyotype.

### 3.2 Donor 2395 iPSC differentiate into insulin producing beta-like cells that respond to glucose

β-cells participate in the pathogenesis of T1D by shedding antigen (Figure 1, Step1) to initiate the autoimmune response and are also targeted for destruction by the immune system in T1D (Figure 1, Step 4A). Any fully isogenic system to study these steps in T1D (Figure 1) will require a renewable source of islets or β-cells. Stem cell-derived β-cells (sBC) can fulfill this need. With published protocols for differentiation of β-cells available, we proceeded to differentiate iPSC from the three lines (1-018, 1-023, and 2395) into sBC. Lines 1-018 and 1-023 have previously been differentiated into validated sBC in islet like clusters with expression of c-peptide and secretion of insulin (25, 46). iPSC from line 2395 were differentiated to sBC by first differentiating to definitive endoderm (DE). Cultures were only moved forward if they reached >80% SOX17^+^ at the DE stage (Figure 3A). At the β-like stage, cells reached 37.8 ± 3.5% insulin^+^glucagon^-^ (Figure 3 B & C; n = 3). Fluorescent micrographs (Figure 3D) demonstrate an islet-like cluster of sBC from 2395 with cells expressing NXK6.1, PDX-1, and c-peptide. To verify that the sBC were functional, those differentiated from 1-018, 1-023, and 2395 were subjected to static glucose-stimulated insulin secretory assays and compared to primary human islets. Clusters from all three lines secreted insulin, albeit less than the average stimulation index of primary islets (Figure 3E). While there were no significant differences, 1-023 and 2395 out-performed clusters from 1-018 on average. In summary these lines differentiated into glucose-responsive sBC allowing a first step towards building *in vitro* systems for studying the interactions of isogenic immune cells with β-cells.

**Figure 3.**
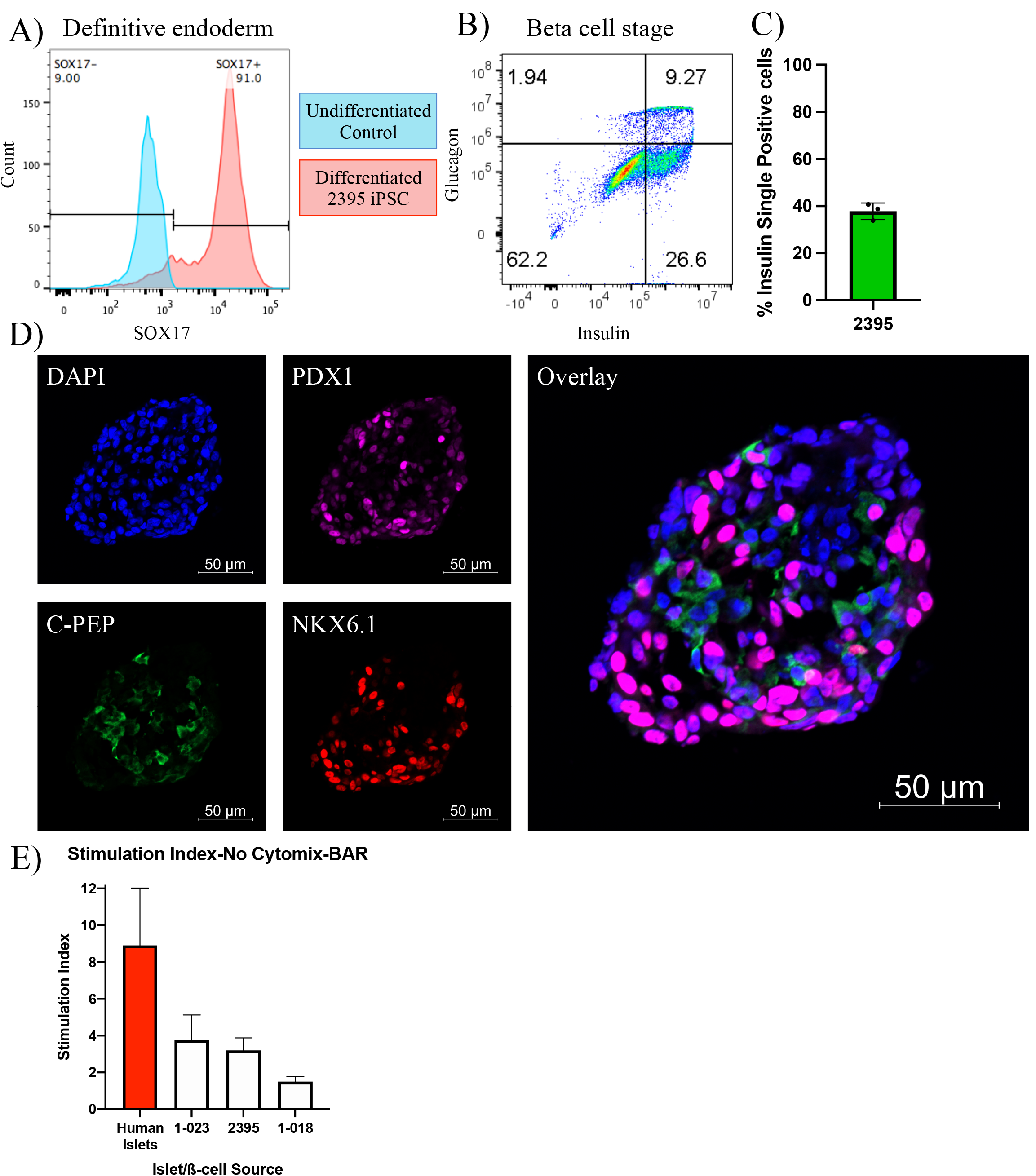
Donor 2395 iPSC differentiate to stem cell derived beta-like cells (sBC). **(A)** Donor 2395 iPSC upregulate SOX17 when differentiated to definitive endoderm. **(B)** When donor 2395 definitive endoderm is differentiated the β-cell stage, many polyhormonal (insulin^+^glucagon^+^) and monohormonal (insulin^+^) cells arise. **(C)** The percent of insulin^+^glucagon^-^ cells from 3 repeats of 2395 sBC differentiation. **(D)** An sBC from donor 2395 was fixed, labeled with DAPI, stained with antibodies against PDX1, C-peptide, and NKX6.1 revealing many C-peptide/NKX6.1 double positive β-cells. **(E)** Donor 2395 sBC as well as iPS-β-cells from lines 1-018 and 1-023 exhibit glucose-stimulated insulin secretion. Stimulation index = insulin secretion in 16.7mM glucose/insulin secretion in 3mM glucose.

### 3.3 HLA Typing of iPSC 1-018, 1-023, and 2395

Class II HLA alleles confer the strongest genetic risk for pathogenesis of T1D, highlighting the importance of CD4^+^ T cells. CD8^+^ T cells are the predominant cell in human insulitis emphasizing that HLA Class I alleles play a major role in T1D development. HLA Class I regulates many of the key steps thought to be vital for T1D initiation such as T cell activation and targeted deletion of pancreatic β-cells (Figure 1). For these reasons it is essential that iPSC harbor useful HLA alleles for the study of T1D. HLA alleles were determined for all three iPSC lines (Table 1). The studies here will focus on the interactions of HLA-A*0201 restricted CD8^+^ T cell avatars (20) with iPSC-derived cells expressing HLA-A*0201 with cognate peptide. Lines 2395 and 1-023 carry HLA-A*0201 while 1-018 does not (Table 1). Therefore, we prioritized the former two iPSC lines for further studies.

**Table.**
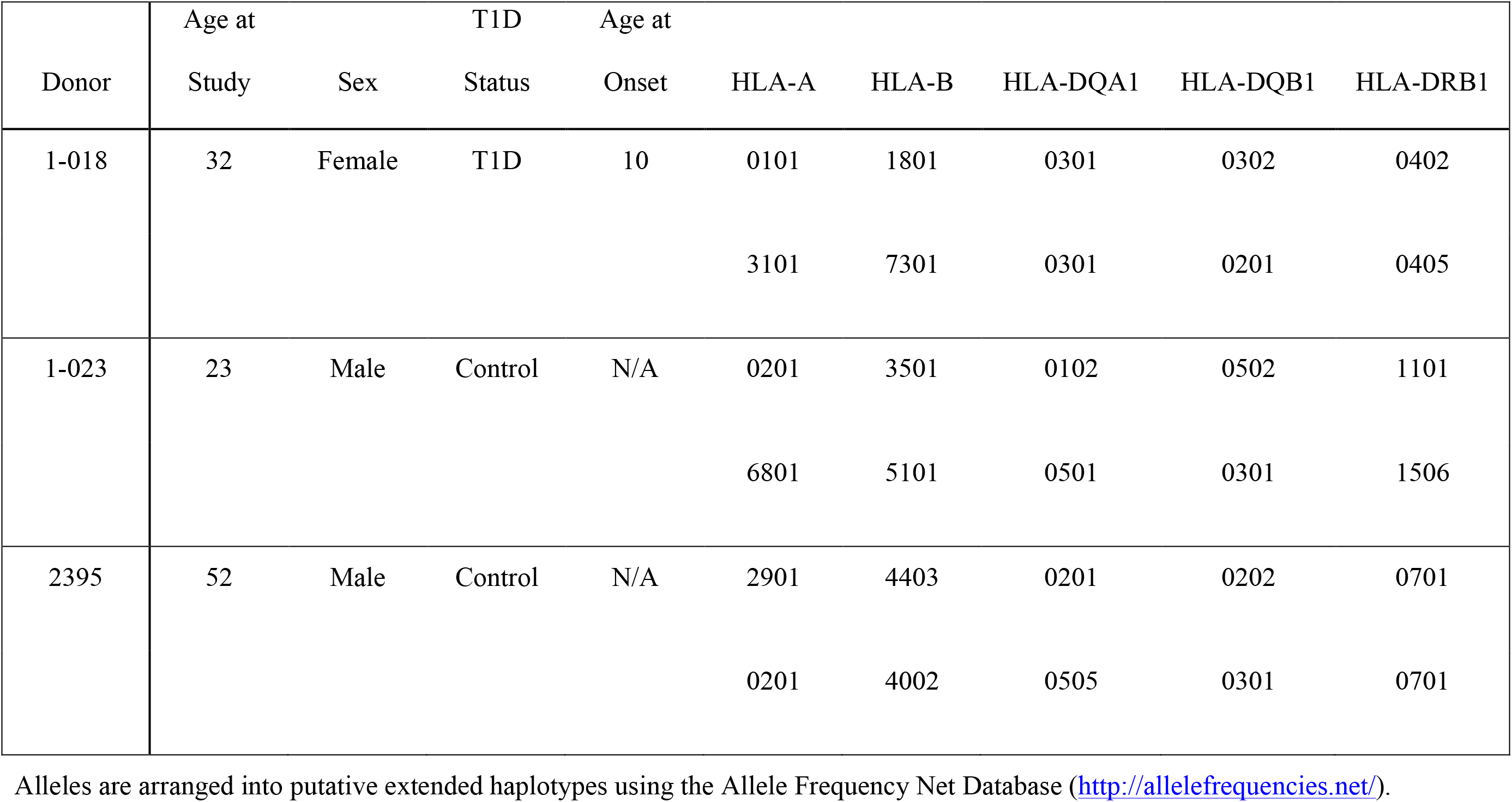
iPSC Donor Demographics and HLA Alleles for lines 1-018, 1-023, and 2395.

| Donor | Age at<br>Study | Sex | T1D<br>Status | Age at<br>Onset | HLA-A | HLA-B | HLA-DQA1 | HLA-DQB1 | HLA-DRB1 |
| --- | --- | --- | --- | --- | --- | --- | --- | --- | --- |
| 1-018 | 32 | Female | T1D | 10 | 0101 | 1801 | 0301 | 0302 | 0402 |
|  |  |  |  |  | 3101 | 7301 | 0301 | 0201 | 0405 |
| 1-023 | 23 | Male | Control | N/A | 0201 | 3501 | 0102 | 0502 | 1101 |
|  |  |  |  |  | 6801 | 5101 | 0501 | 0301 | 1506 |
| 2395 | 52 | Male | Control | N/A | 2901 | 4403 | 0201 | 0202 | 0701 |
|  |  |  |  |  | 0201 | 4002 | 0505 | 0301 | 0701 |

Alleles are arranged into putative extended haplotypes using the Allele Frequency Net Database (http://allelefrequencies.net/).

### 3.4 Donor 2395 iPSC Efficiently Differentiate to Monocytes, MDM, and MoDC

While the initiating events in T1D are not decisively characterized, initiating a productive T cell response requires at least two signals during the interaction of APC with T cells (Figure 1, Step 2). The first signal is binding of the TCR on the T cell to its cognate antigen/MHC complex on an APC and the second signal is the costimulatory signal in the form of co-receptors on the T cell binding their respective ligands on an APC (e.g. CD28 binding CD80/CD86, CD27 binding CD70, CD226 binding CD155, OX-40 binding OX-40L, etc.) (47, 48). APC such as DC and macrophages serve to initiate T cell responses in the lymph nodes and enhance T cell responses at sites of inflammation. In order to model interactions between macrophages and T cells or DC and T cells, we differentiated donor 2395 iPSC to monocytes, then MDM and moDC (Figure 4). Similar to primary classical peripheral blood monocytes, these iPSC-monocytes are CD14^+^CD64^+^CD16^-^ (Figure 4A). Also similar to their primary counterparts, iPSC-MDM are CD11b^+^CD68^+^ (Figure 4B) and iPSC-moDC are MHC-II^+^CDllc^+^ (Figure 4C). Attempts were made to differentiate the iPSC line, 1-023, to monocytes however they never produced CD34^+^ hematopoietic progenitors following hematopoietic differentiation and they also never produced CD14^+^ monocytes following monocyte induction. Due to the inability of 1-023 iPSC to differentiate into Monocytes, MDM, and MoDC we prioritized all iPSC 2395 for the remaining studies.

**Figure 4.**
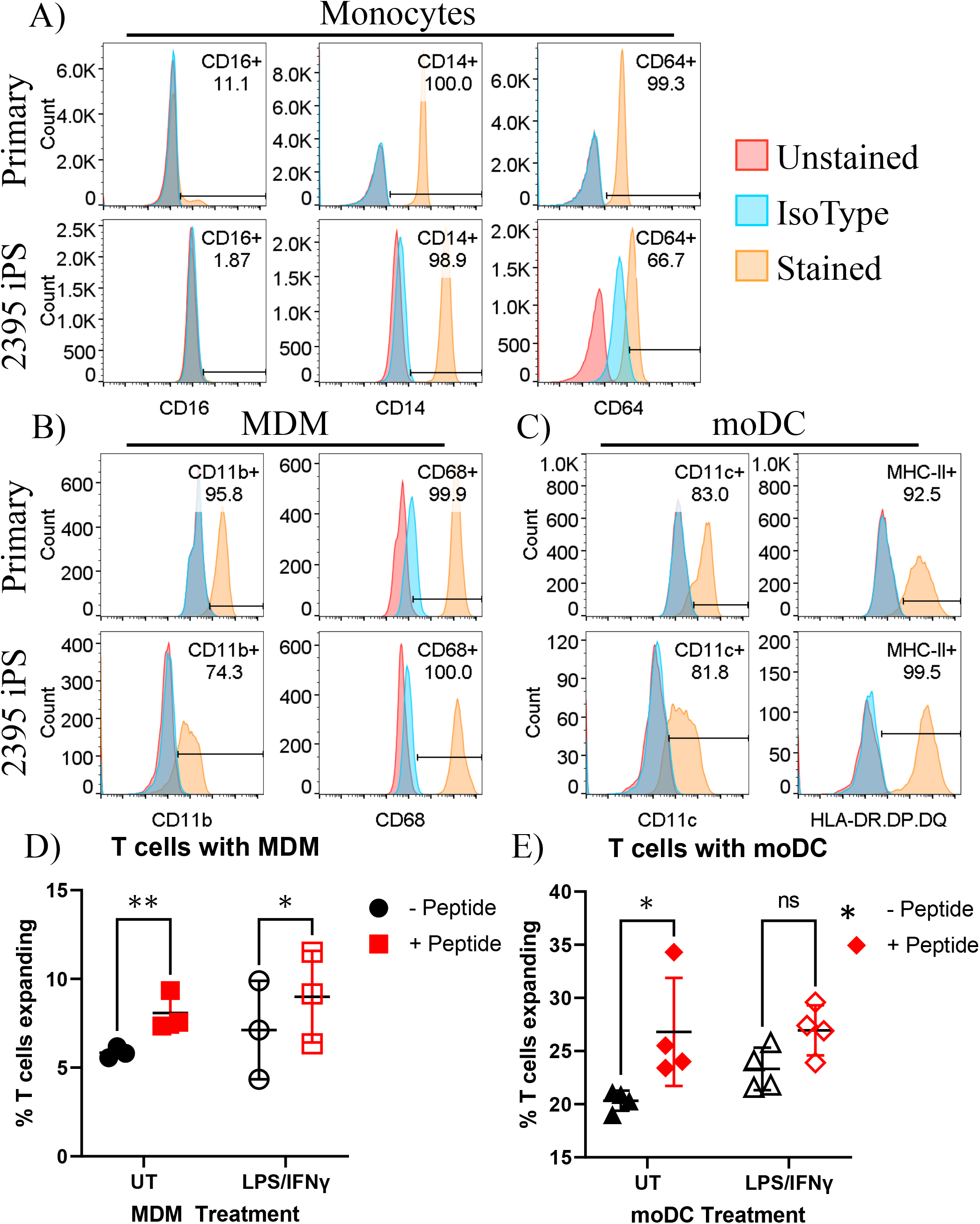
Donor 2395 iPSC differentiate efficiently to monocytes, monocyte-derived macrophages (MDM) and monocyte-derived dendritic cells (moDC) and induce antigen specific CD4^+^ and CD8^+^ T cell expansion **(A)** Donor 2395 iPSC-derived monocytes are CD16^-^, CD14^+^, and CD64^+^ similar to primary classical monocytes isolated from blood. **(B)** Donor 2395 iPSC-derived MDM express CD11b and CD68 similar to primary MDM. **(C)** Donor 2395 iPSC-derived moDC express CD11c and MHC-II similar to primary moDC. **(D)** iPSC-MDM expand Total T cells in an antigen specific manner. When antigen (CEFX peptide pools) is present, there is a significantly increased expansion of T cells compared to when Total T cells are cultured with APC alone (ANOVAs **p = 0.0011 with n=3). **(E)** iPSC-moDC expand Total T cells in an antigen specific manner. When antigen is present, there is a significantly increased expansion of T cells compared to when Total T cells are cultured with APC alone (ANOVAs *p = 0.0104 with n=4).

### 3.5 Donor 2395 iPSC-moDC and iPSC-MDM Elicit Donor 2395 CD4^+^ and CD8^+^ T Cell Expansion in an Antigen Specific Manner

To determine if iPSC-moDC and MDM are able to elicit an antigen specific CD4^+^ and CD8^+^ T cell response, iPSC-moDC or MDM from donor 2395 were activated with LPS/IFNγ, loaded with CEFX Ultra SuperStim Pool MHC-I and MHC-II subsets, and co-cultured with CFSE (ThermoFisher) stained total T cells from donor 2395. When 2395 total T cells were cultured with iPSC-MDM alone (Figure 4D; closed black circles), or LPS/IFNγ-activated iPSC-MDM (Figure 4D; open black circles) there was little T cell expansion observed. However, when 2395 total T cells were cultured with CEFX peptide-loaded MDM (Figure 4D; closed red squares), or LPS/IFNγ-treated, CEFX peptide-loaded iPSC-MDM (Figure 4D; open red squares), there was significantly more expansion than when they were cultured with iPSC-MDM without peptide loading. iPSC-moDC also induced T cell expansion in an antigen specific manner. When total T cells were cultured with peptide-loaded iPSC-moDC (Figure 4E; closed red diamonds), expansion was increased compared to T cells cultured with iPSC-moDC alone (Figure 4E; closed black triangles). However, when iPSC-moDC were activated with LPS/IFNγ first, there was no significant difference in T cell expansion when iPSC-moDC were peptide loaded (Figure 4E; open red diamond) compared to iPSC-moDC without peptide loading (Figure 4E; open black triangles).

### 3.6 Donor 2395 CD8^+^ T Cell Avatars Bind to Donor 2395 EC in an Antigen Specific Manner

After T cells are activated, they must traffic to the islets of Langerhans. To exit the islet capillaries and enter the islets, T cells must migrate across the endothelial cell (EC) layer lining the insides of the capillaries in a process called extravasation [(49) and Figure 1, Step 3]. In this process, T cells first migrate to the site of inflammation via chemotaxis, they then must tether and roll across activated ECs before adhering, and finally transmigrating across the epithelium (49). To model the interactions of CD8^+^ T cells with vascular endothelium, we assessed CD8^+^ T cell avatars adhesion to an antigen-loaded iEC monolayer using cells from the same donor. As described recently (38), 2395 iPSC efficiently differentiate into iEC with morphology, cell surface marker profile, and functional capabilities similar to human aortic endothelial cells (HAEC). Here an *in vitro* hydrodynamic flow chamber adhesion assay was performed with monolayers of iEC from donor 2395 and either IGRP avatars or MART-1 avatars from 2395. When the 2395 iEC monolayer was not presenting antigenic peptide, the 2395 IGRP avatars failed to adhere strongly and were displaced at higher laminar flow rates (Figure 5A; black). Similarly, when the 2395 iEC monolayer was pulsed with the peptide antigen recognized by the MART-1 TCR, Melan-A_26-35_ (EAAGIGILTV), 2395 IGRP avatars detached at higher flow rates (Figure 5A; red). In contrast, 2395 IGRP avatars formed strong adhesions with the IGRP_265-273_ peptide-pulsed iEC monolayer and remained attached at higher flow rates (Figure 5A; gray). We confirmed the system’s utility with a second TCR and antigen, 2395 MART-1 avatars and the peptide it recognizes, Melan-A_26-35_. Using the laminar flow system without peptide the MART-1 avatars failed to form firm adhesions with the iEC monolayer (Figure 5B; black). Addition of IGRP_265-273_ did not facilitate adhesion as this peptide is not recognized by the MART-1 avatars (Figure 5A; gray). In contrast, 2395 MART-1 avatars made firm adhesions with the Melan-A_26-35_ peptide-pulsed iEC monolayer as observed by binding at higher flow rates (Figure 5A; red). As vascular adhesion interactions are critical for T cell trafficking into pancreatic islets for targeted destruction of β-cells in T1D (42,50–52), this proof-of-concept study demonstrates the utility of an antigen-dependent system for assessing interactions of T cells with endothelial cells. The ability to model these interactions with human cells allows for intensive interrogation of genetics regulating vascular inflammation or therapeutics meant to target the disruption of lymphocyte adhesion to vascular cells (Figure 1, Step 3).

**Figure 5.**
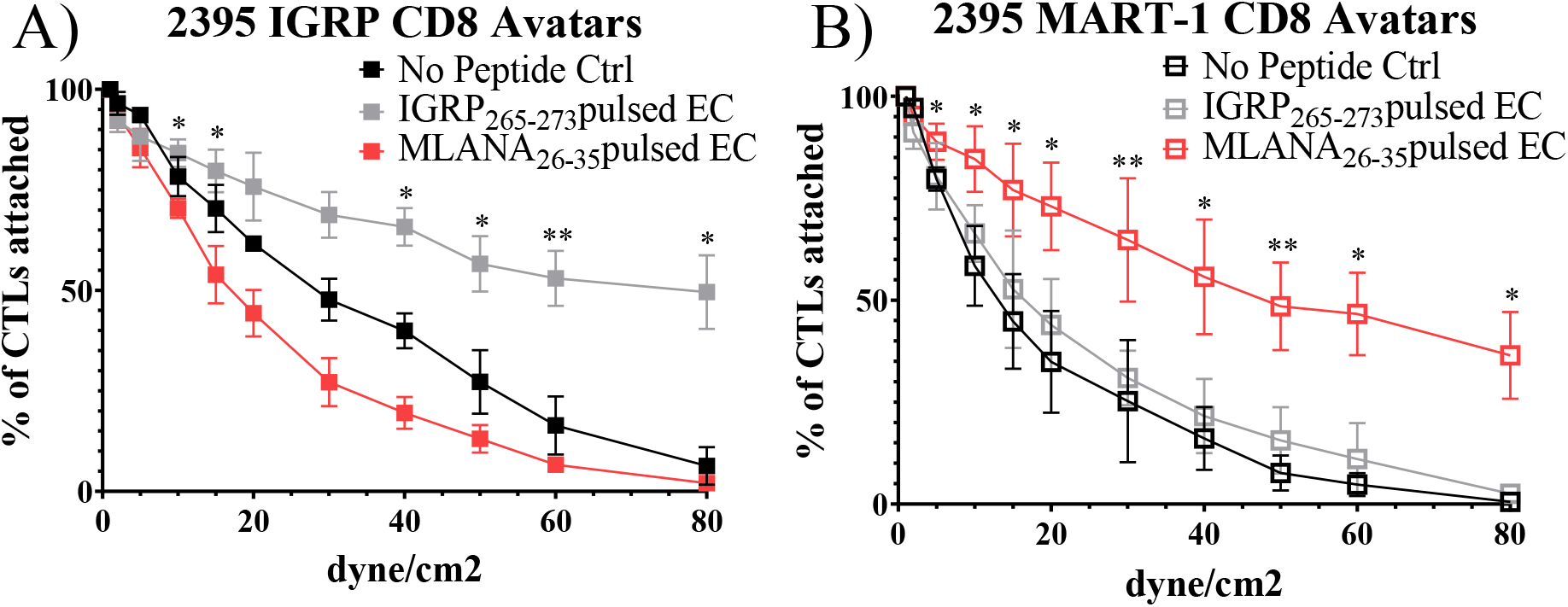
Donor 2395 CD8^+^ T cell avatars exhibit antigen specific binding to donor 2395 iPS-EC. **(A)** When donor 2395 IGRP CD8^+^ T cell avatars are flowed across a donor 2395 iPSC-derived endothelial cell (iPS-EC) monolayer loaded with IGRP peptide, the IGRP CD8^+^ T cell avatars adhere more than MART-1 CD8^+^ T cell avatars or when the iPS-EC is not loaded with peptide. **(B)** When donor 2395 MART-1 CD8^+^ T cell avatars are flowed across a donor 2395 iPSC-derived endothelial cell (iPS-EC) monolayer loaded with Melan-A_26-35_, the MART-1 CD8^+^ T cell avatars adhere more than IGRP CD8^+^ T cell avatars or when the iPS-EC is not loaded with peptide. n=3 for all *in vitro* hydrodynamic flow assay experiments. * indicates that p < 0.05 and ** indicates that p < 0.01 in 2-way ANOVA with multiple comparisons.

### 3.7 Donor 2395 CD8^+^ T Cell Avatars Efficiently Target Donor 2395 sBC in an Antigen Specific Manner

To determine if sBC are able to be targeted by CD8^+^ T cells in an antigen specific manner, a cell mediated lymphocytocixity (CML) assay was run with dispersed clusters of sBC differentiated from 1-023, 2395, or 1-018 as targets. The HLA-A*0201^+^ human βL5 cell line and dispersed HLA-A*0201^+^ human islet cells were used as positive controls (Figure 6). These groups were co-cultured with donor 2395 T cell avatars expressing a T cell receptor that recognizes a peptide from IGRP in the context of HLA-A*0201 (40, 41). The 2395 IGRP-avatars were able to target and kill the HLA-A*0201 expressing βL5 and primary human islets (Figure 6A). In addition, cells differentiated from 1-023 and 2395 were lysed by the avatars (Figure 6A) as they carry HLA-A*0201 (Table 1). Cells from 1-018 were not killed by the avatars (Figure 6A) as these cells do not have the correct HLA restriction (Table 1). We also assessed the ability of 2395 MART-1 T cell avatars that recognize a non-β-cell peptide derived from Melan-A_26-35_ (20, 53) to target and destroy cells that were pulsed with this Melan-A peptide or left untreated. MART-1-avatars were able to target and lyse both βL5 and 2395 sBC when the peptide was present in a fashion that was regulated by the number of MART-1 avatars added (Figure 6B). However, when the βL5 or 2395 sBC were not pulsed with Melan-A_26-35_, neither population of cells was lysed. These results indicate that sBC can be utilized to study the interactions of β-cells with antigen specific T cells. In addition, as 2395 T cell avatars kill 2395 sBC in an antigen-specific fashion (Figure 6B) this isogenic system is a viable method to study interactions of autoreactive T cells with β-cells.

**Figure 6.**
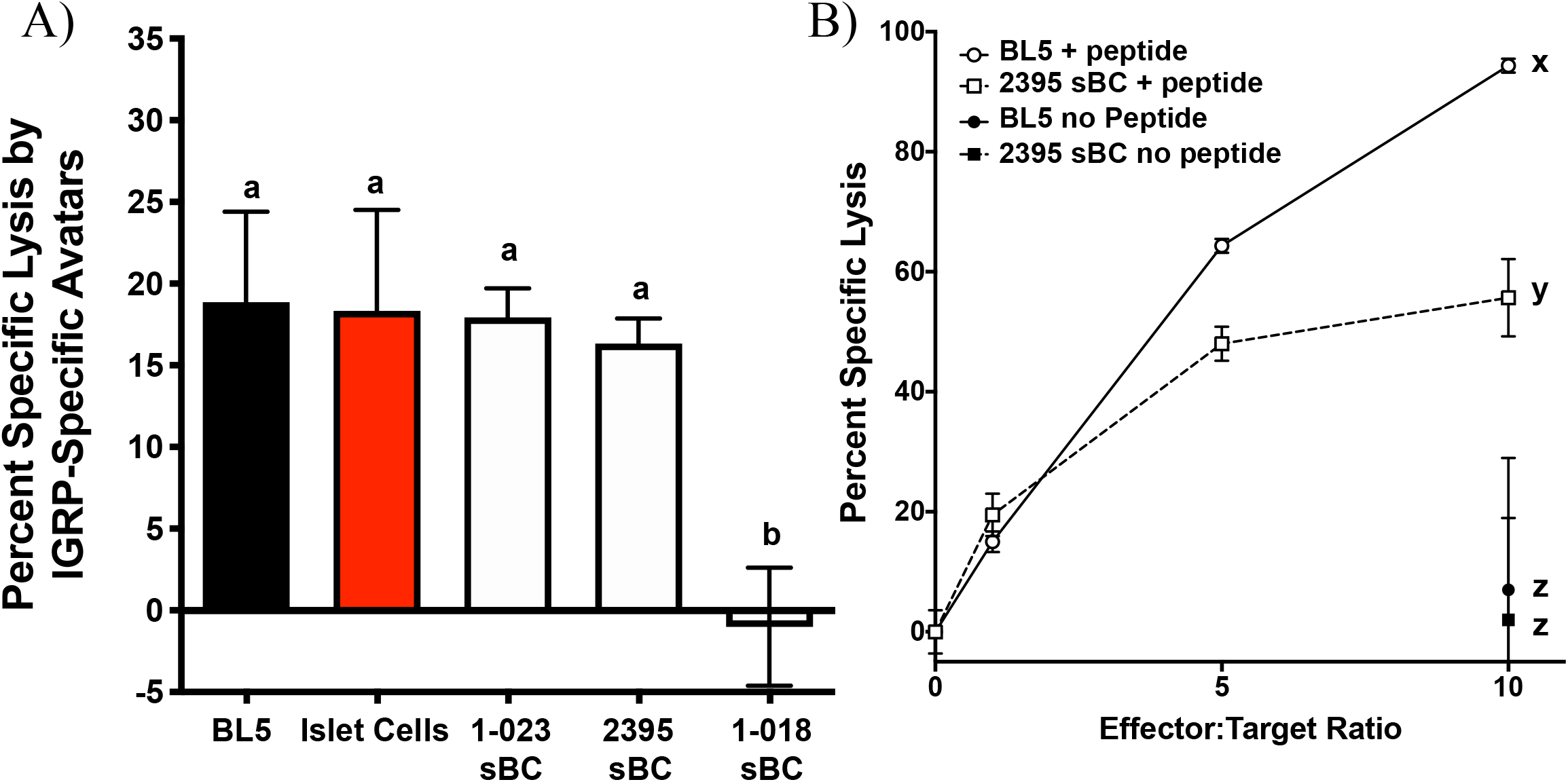
sBC are efficiently targeted by donor 2395 CD8^+^ T cell avatars in an antigen specific and HLA restricted manner. (**A**) Donor 2395 IGPR T cell avatars were co-cultured with βL5 cells, primary islet cells from a HLA-A*0201 positive donor, 1-023 sBC, or 2395 sBC the cells were effectively lysed at an effector to target (E:T) ratio of 10:1. As 1-018 sBC do not harbor HLA-A*0201 these cells were not lysed by these T cell avatars as they require the IGRP peptide to be presented by HLA-A*0201. (**B**) Donor 2395 MART-1-avatars were co-cultured with either βL5 cells or donor 2395 sBC at increasing E:T ratios (0:1, 1:1, 5:1, and 10:1). When pulsed with Melan-A_26-35_, lysis of both βL5 and 2395 sBC increased with escalating E:T ratios. In the absence of Melan-A_26-35_, no lysis was measured. *Letters denote significance: groups/lines with different letters were statistically significant (P<0.05), whereas those with the same letter were not statistically different. These data are from at least 3 independent studies performed in triplicate.

## 4 Discussion and Conclusion

While T1D develops in genetically at-risk individuals, our current understanding of the pathogenesis of T1D in humans continues to evolve due to the complexities of genetic risk in combination with environmental factors (54). Recent GWAS efforts have dramatically expanded the number of loci that are associated with both increased or decreased risk for T1D (1,2,11,12,3–10). The mechanisms linking the majority of these polymorphisms, alleles, and allotypes to the development of T1D are not clear. Disease heterogeneity in T1D may result from the dissimilar combinations of genes inherited by at risk individuals that instigate variations in onset age and disease endotypes. Indeed, specific combinations of risk HLA haplotypes have been linked to earlier age of onset as well as changes in the immune cell components of the insulitic lesion (55). Due to the impact of total combined genetic risk, it is necessary to sufficiently power mechanistic studies with large sample sizes to determine the responsibility of a single polymorphism to T1D and the cell types influenced by the greater than 100 polymorphisms that are linked to T1D (1,2,11,12,3–10). With the development of isogenic systems, such as that described here, methodologies have advanced that allow for interrogation of the effect of a single polymorphism in the absence of other genetic differences. Models derived from primary cells can lack reproducibility because certain cell types (i.e. β-cells) are not renewable; utilizing iPSC circumvents this issue. Herein we have shown that key cell types involved in the pathogenesis of T1D (Figure 1) can be derived from a single donor and used to construct an isogenic system to interrogate relevant cell-cell interactions. This renewable system provides a powerful platform to interrogate how specific genetic polymorphisms influence both cell function and cell-cell interactions in isolation of other genetic differences.

Thus far, we have derived monocytes, macrophages, dendritic cells (DC), endothelial cells (EC), beta-like cells (sBC), and CD8^+^ T cell avatars all from donor 2395. In this isogenic system derived from donor 2395, we have shown that both macrophages and DC can expand total T cells (Figure 4), that CD8^+^ T cell avatars bind to EC in an antigen specific manner (Figure 5), and that CD8^+^ T cell avatars can target and lyse sBC that is dependent on the trimolecular complex (Figure 6). The current system affords power for interrogating these simple cell-cell interactions as well as more complex cell-cell interactions such as macrophage, T cell, and β-cell interactions. While the system has great utility, it does however have limitations.

One of the current requirements of the current system is the need to recall specific donors to obtain T cells. This necessitates proper IRB approval and flexibility of research participants and investigators to schedule blood draws. Currently there are protocols for obtaining T cells from iPSC however these produce predominantly CD8^+^ T cells and very few CD4^+^ T cells (56, 57). The addition of iPSC-derived naïve autoreactive CD8^+^ T cells to this system can be implemented to provide yet another way for interrogation of cell-cell interactions and eliminate the need to recall donors to obtain CD8^+^ T cells. iPSC can be transduced with lentiviral vectors (58, 59), therefore a system to develop antigen specific iPSC-T cells could be employed wherein the iPSC are engineered to express a specific TCR to recognize self or nominal antigens.

Another shortcoming of this system and iPSC in general is that not all iPSC lines efficiently differentiate into all necessary cell types. For example, 1-018 sBC were not as functional as sBC from lines 1-023 or 2395 (Figure 3E). In addition, while 1-023 differentiated well into sBC, this iPSC line did not produce monocytes. Lack of monocytes prevented downstream differentiation to macrophages and dendritic cells. Other groups have observed that lineage differentiation capacity is dependent on the iPSC line necessitating protocol optimization for lines that fail to differentiate effectively (60). As such, methodological modifications specific for 1-023 may resolve the observed dearth of monocytes from this line. However, the necessity of individualized differentiation protocols could impact the throughput of any system. Another important consideration and potential difficulty results from the complexity of T1D genetics. This requires careful consideration of the cells utilized for differentiation and study. Due to the strong genetic influence of the HLA super locus on T1D and the need for specific HLA alleles to be present for antigen specific studies, genotyping and selecting iPSC with HLA alleles (Table 1) that match TCRs to be utilized is essential. Here we eliminated 1-018 from the system as this line did not have the necessary HLA-A*0201 allele to interact with the T cell avatars (Table 1). Indeed, 1-018 sBC were not targeted and lysed by the IGRP-Avatars (Figure 6A). The HLA Class I and Class II loci are the most polymorphic coding genes in the human genome (61), therefore matching HLA for the current study was selected. However, advances in gene editing technology may, in future studies, allow for extensive modification of HLA loci and obviate the requirement for excluding cells based on HLA allele types. Regardless of the more difficult requirements of the current system, there are many developing technologies that will improve future iterations.

One of the currently developing technologies is gene editing in iPSC. Gene editing can increase the flexibility of this system. There are numerous protocols for editing SNPs or knocking out genes in iPSC. While knockout is straightforward and there are high-efficiency protocols available for iPSC, easy to implement and high-efficiency protocols for scarless editing of single bases, such as SNPs, are still being developed (62–68). Once these high-efficiency protocols are developed, editing SNPs in iPSC will allow us to interrogate how credible risk or resistance SNPs influence these cell-cell interactions. The ability to edit iPSC donor lines to harbor both the T1D-risk and T1D-resistance alleles at a single locus will potentially suppress variation by allowing edited donor lines to be compared to each other. These methods would have exceptional value to study the genetic influences in cells, organoids, or organs that are difficult to access or genetically modify in a homogenous fashion. This would greatly facilitate the study of rare genetic variants (e.g. *rs34536443* in *TYK2*, *rs2476601* in *PTPN22*, *rs35744605* in *IFIH1*, etc.) as editing iPSC lines to the rare minor alleles at T1D risk loci would allow for generation of numerous lines from different donors that could be studied in a pairwise fashion versus identifying and specifically recruiting donors with minor allele frequency (MAF) <0.05. For example, the frequency of individuals homozygous for the rare allele of *rs34536443* in *TYK2* was recently reported to be 213 per 100,000 with MAF =0.04 (69, 70). The ability to edit a credible risk or resistance SNP will facilitate determination of the impact of these rare variants and likely bolster efforts seeking to resolve the actual SNP(s) in linkage disequilibrium (LD) that influence disease risk (e.g. the 233kb LD block on chromosome 16p13 that contains *CLEC16A*) (71–73). In addition, other technologies in development, such as ESC and iPSC-derived thymii that can output functional naïve T cells, are novel tools for studying human disease etiology. As thymic epithelial cells (iPSC-TEC) (74, 75) have the potential to regulate T cell production from progenitors, use of these evolving technologies in combination (gene editing and iPSC-TEC) would allow for studies to dissect how T1D-linked loci, or those linked to other diseases, regulate T cell development. These and other currently advancing technologies will greatly enhance isogenic systems, improving their capacity to model human diseases.

Isogenic systems derived from iPSC will greatly augment our ability to interrogate key cellular interactions involved in genetic regulation of T1D pathogenesis. While the system built here is focused on T1D, it can be utilized to study cell-cell interactions and genetic regulation in the context of other human diseases. In summary, we have presented an isogenic model that can currently be used to interrogate many T1D-relevant cell-cell interactions. As more technologies for gene editing and differentiating iPSC continue to be developed these tools can be implemented to build a more relevant model capable of answering more complex questions.

## 5 Author Contributions

LHA, SES, CEM, and MAW conceived the idea, designed the research plan, and participated in all aspects of writing the manuscript. KES, LS, SOO, BNN, MKH, JPT, NF, PS, IR, RC-G, DJP, HR, AS, NT, TMB, JC, DE performed experiments, analyzed data, interpreted results of experiments, and edited the manuscript. All authors contributed to the article and approved the submitted version.

## 6 Acknowledgments

The University of Florida Center for Immunology and Transplantation, The University of Florida Center for Cellular Reprogramming, The University of Florida Diabetes Institute, The University of Florida Clinical and Translational Science Institute, and the NIH funded Integrated Islet Distribution Program were all essential to the completion of these studies. This intra-institutional collaboration was facilitated by the Human Islet Research Network (https://hirnetwork.org/).

---

**Supplemental Table.** Antibody clones and catalog numbers.

| Antigen | Fluorophore | Source | Clone | Assay | Company | Catalog # |
| --- | --- | --- | --- | --- | --- | --- |
| OCT4 | PE | mouse | 3A2A20 | FC | BioLegend | 653704 |
| SSEA-4 | APC | mouse | MC-813-70 | FC | BioLegend | 330418 |
| SSEA-1 | PE | mouse | HI98 | FC | BioLegend | 301906 |
| Nestin | PE | mouse | 25/NESTIN | FC | BD | 561230 |
| FoxA2 | PE | mouse | N17-280 | FC | BD | 561589 |
| Sox17 | AF488 | mouse | P7-969 | FC | BD | 562205 |
| NCAM | PE | mouse | HCD56 | FC | StemCell | 60021PE.1 |
| Brachyury | APC | goat | polyclonal | FC | R&D | IC2085A |
| NKX6.1 | PE | mouse | R11-560 | FC | BD | 563023 |
| PDX1 | AF488 | mouse | 658A5 | FC | BD | 562274 |
| Glucagon | BV421 | mouse | U16-850 | FC | BD | 565891 |
| C-peptide | AF647 | mouse | U8-424 | FC | BD | 565831 |
| PDX1 | n/a | goat | polyclonal | IF | R&D | AF2419 |
| anti-Goat | AF647 | donkey | polyclonal | IF | Thermo | A-21447 |
| C-peptide | n/a | rat | n/a | IF | DSHB | GN-ID4 |
| anti-Rat | AF488 | donkey | polyclonal | IF | Thermo | A-21208 |
| NKX6.1 | n/a | mouse | n/a | IF | DSHB | F55A10 |
| anti-mouse | AF555 | donkey | polyclonal | IF | Thermo | A-31570 |
| CD14 | PE | mouse | M5E2 | FC | BioLegend | 301806 |
| CD16 | FITC | mouse | eBioCB16 | FC | ThermoFisher | 11-0168-42 |
| CD64 | APC | mouse | 10.1 | FC | ThermoFisher | 17-0649-42 |
| CD11b | FITC | mouse | ICRF44 | FC | BioLegend | 301330 |
| CD68 | PE | mouse | eBioY1/82A | FC | ThermoFisher | 12-0689-41 |
| CD11c | APC | mouse | Bu15 | FC | BioLegend | 337208 |
| HLA-DR, DP, DQ | APC | mouse | Tü39 | FC | BioLegend | 361714 |

\*FC = Flow Cytometry, IF = Immunofluorescence

